# Haplotype rather than single causal variants effects contribute to regulatory gene expression associations in human myeloid cells

**DOI:** 10.1101/2025.01.30.635675

**Authors:** Emily Greenwood, Mingming Cao, Ciaran M. Lee, Aidi Liu, Buhle Moyo, Gang Bao, Greg Gibson

**Author notes:** Joint corresponding authors. These authors contributed equally to this work (Emily Greenwood, Mingming Cao).

## Abstract

Genome-wide association studies typically identify hundreds to thousands of loci, many of which harbor multiple independent peaks, each parsimoniously assumed to be due to the activity of a single causal variant. Fine-mapping of such variants has become a priority and since most associations are located within regulatory regions, it is also assumed that they colocalize with regulatory variants that influence the expression of nearby genes. Here we examine these assumptions by using a moderate throughput expression CROPseq protocol in which Cas9 nuclease is used to induce small insertions and deletions across the credible set of SNPs that may account for expression quantitative trait loci (eQTL) for genes associated with inflammatory bowel disease (IBD). Of the 4,384 SNPs targeted in 88 loci (an average of 50 per locus), 439 were significant and further examined for validation. From these, 98 significantly altered target gene expression in HL-60 myeloid cell line, 74 in induced macrophages from these HL-60 cells, and 78 in induced neutrophils for a total of 201 validated effects (46%), 43 of which were observed in at least two of the cell types. Considering the observed sensitivity and specificity of the controls, we estimate that there are at least 150 true positives per cell type, an average of almost 2.4 for each of the 64 eQTL for which putative causal variants have been fine-mapped. This implies that haplotype effects are likely to explain many of the associations. We also demonstrate that the same approach can be used to investigate the activity of very rare variants in regulatory regions for 89 genes, providing a rapid strategy for establishing clinical relevance of non-coding mutations.

## BACKGROUND

Genome-wide association studies (GWAS) have revolutionized geneticists’ ability to identify loci that influence variance in complex traits^1–3^. Almost all common polygenic contributions to height, for example, map to 12,111 loci distributed among one fifth of human genes^4^, implying that multiple variants influence the activity of each gene^5^. Studies with tens of thousands of cases and controls typically uncover several hundred loci contributing to disease liability, as typified by inflammatory bowel disease (IBD) for which 250 loci have been reported^6,7^. However, only a small minority of these associations have been fine-mapped to causal variants, many involving amino acid encoding variants^8–10^. Regulatory complexity seems to be much greater yet is responsible for over ninety percent of effects and requires experimental strategies to resolve single nucleotide contributions.

Two broad strategies for regulatory fine-mapping have emerged: CRISPR screenings and massively parallel reporter assays (MPRA). CRISPR interference utilizes deactivated Cas9 nuclease (dCas9) coupled to a KRAB domain to repress gene expression mediated by suspected enhancers that are targeted by a panel of guide RNAs (gRNAs). It has efficiently demonstrated that peak eQTL variants located in open chromatin regions of relevant cell types are strongly enriched for regulatory activity^11,12^. For example, STING-seq^13^ provided variant-to-function linkage of blood-cell trait GWAS polymorphisms for 134 cis-regulatory elements to the closest gene’s expression. MPRAs, by contrast, clone random DNA fragments adjacent to a reporter gene, the expression of which is used to identify fragments that harbor enhancer activity^14,15^. Contrasting activity of fragments containing either allele at a specific site provides prima facie evidence for regulatory polymorphism, one study covering almost half the genome revealing over 30,000 potentially causal variants that only partially overlapped in two cell types^16^.

Both strategies have provided evidence that single GWAS signals should sometimes be attributed to more than one SNP, but this proposition has not been systematically evaluated. While it is parsimonious to assume a one-to-one mapping of statistical signal to causal variant, a reasonable alternative possibility is that haplotype effects are common, where two or more variants in high linkage disequilibrium (LD) are responsible for a given signal^17^. Evidence of this would have broad implications for understanding the genetic architecture of complex traits, resolving the impacts of soft selection and genetic drift on evolutionary dynamics^18–20^, and interpreting the transferability of polygenic scores within and across ancestry groups^21–23^. Here we adapt CROPseq^24^, a CRISPR/Cas9 based strategy for moderate parallel genome editing, to fine-map causal regulatory variants, demonstrating that 66 eQTL signals for 52 IBD GWAS implicated genes are indeed often attributable to multiple SNPs in the HL-60 myeloid cell line and derivatives. Thus, regulatory complexity includes multiple independent signals for each gene as well as multiple variants contributing to each signal.

## RESULTS

### Expression Crop-Seq Mapping of Rare Variants

Our experimental approach is to target credible regulatory polymorphisms for genome editing in the form of mostly small deletions in single cells of the HL-60 myeloid cell line engineered to constitutively express Cas9. A pool of regulatory variant specific gRNAs is introduced and expression of the nearby target transcript in edited cells is measured by single-cell RNAseq (scRNAseq). We previously illustrated this approach^25^ by fine-mapping causal variants at eQTL peaks for two IBD risk genes, *CISD1* and *PARK7*, for which dose-dependent effects were validated in clonal cell lines.

Gene expression outliers detected in the GTEx Project are often linked to rare variants located either in the gene body or within 10kb of the promoter^26,27^. To evaluate the ability of expression CROPseq to validate such rare variants (MAF < 1%), we generated a pool of unique gRNAs targeting candidate rare variants previously inferred to affect gene expression in whole blood by Watershed^27^, a probabilistic model of functional annotation. We tested 465 rare candidates for 89 genes, the majority of which were targeted by two distinct gRNAs. Each gene also included rare variants not expected to be functional by Watershed, resulting in 327 negative controls targeted by one gRNA each. Additional positive and negative controls were included to ensure sufficient editing (fig. S1, see Methods). After standard quality-control and removal of cells without a detectable gRNA, 33,302 and 43,129 HL-60 cells in each of the two transductions were analyzed with the Fluent Biosciences platform^28^. At a Bonferroni corrected p-value < 0.001, 5.4X as many Watershed-predicted rare variants disrupted target expression as the negative controls (4.9% versus 0.9%), and strong enrichment of 2.5X was also observed at the more liberal threshold of p-value < 0.05 (34.2% versus 13.8%) (Fig. 1A).

**Figure 1.**
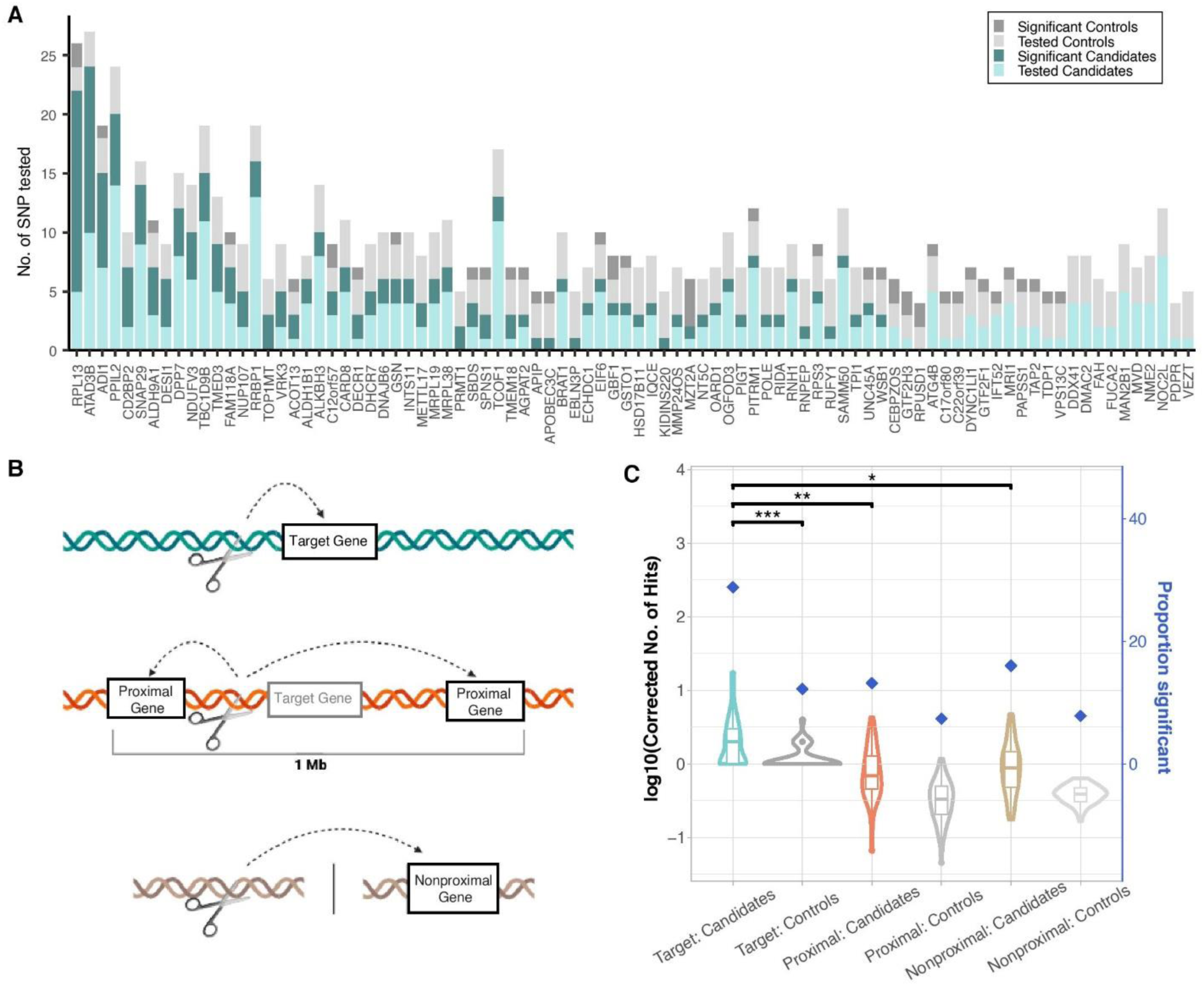
Results of Rare Variant Screening. (A) List of 89 genes tested in rare variant screening encompassing 1,170 gRNA targeting 465 rare candidate variants and 327 controls. Tested but nonsignificant candidates are in light blue (n=306), significant candidates (p-value < 0.05) in dark blue (n=159), tested negative controls in light grey (n=282), and significant negative in dark grey (n=45). (B) Schematic showing the classification of target genes in blue, proximal genes in orange, and nonproximal genes in light brown. Proximal genes are defined as those within a 1Mb region surrounding the target gene and nonproximal genes are outside these regions, elsewhere in the genome. (C) The number of significant hits altering target gene expression, proximal gene expression or nonproximal gene expression for 86 target genes with at least one significant hit. The number of significant candidates (blue) and significant controls (dark grey) altering target gene expression are shown in the first two violin plots. The next two violin plots report the number of significant candidates (orange) and controls (medium grey) altering proximal gene expression. The last two violin plots report the number of significant candidates (tan) and controls (light grey) altering nonproximal gene expression. The number of hits for the proximal and nonproximal genes were corrected by dividing by the total number of proximal and nonproximal genes tested, respectively. The blue diamonds indicate the percentage of variants that were significant for each group, out of all those tested. Denoted comparisons between groups are following Wilcoxon rank sum test with the following ***=0.001, **=0.01, *=0.05.

These results provide evidence that up to 159 of the 465 rare variants are correctly identified as disrupters of gene expression and are likely responsible for the mentioned outlier effects, making expression CROPseq a viable approach to rapid confirmation of clinical significance. The proportion of true positives is approximately as expected given that HL-60 cells represent the myeloid lineage, which is only one third of peripheral blood immune cells. Despite all genes being expressed in HL-60 (fig. S2), the SNP effects may be specific for other cell types. The experiment is also underpowered to detect all variants, since increasing the number of cells receiving each guide increases the likelihood of observing an effect (fig. S3). Exploring the higher-than-expected false positive rate, we additionally asked how often each gRNA also affected expression of proximal genes located within 1Mb of the transcript start site (TSS) for a target gene, or randomly chosen non-proximal genes expressed in HL-60 cells and located elsewhere in the genome (see Methods). Such background effects were approximately one-half as frequently observed for the Watershed predicted variants and the enrichment for proximal and non-proximal effects relative to controls remained similar at 1.8X and 2.1X, respectively (Fig. 1B,C).

### Fine-Mapping Common eQTL within Credible Sets

We then asked whether eCROPseq can identify causal mediators of IBD by fine-mapping eQTL for 87 associated genes^29^ with adequate expression in HL-60 cells (fig. S4). Expression QTL were previously isolated in peripheral blood through all-but-one conditional analysis^30^, where we showed that although most genes have a single signal, some are more complex with multiple overlapping or nonoverlapping signals. For each eQTL, credible sets and surrounding SNPs were targeted in undifferentiated HL-60 cells and the impact on transcript abundance for a total of 4,382 variants was quantified on the 10x Genomics^31^ platform. We detected at least one nominally significant (per gene p-value < 0.005) SNP for 87% of genes, and often observed multiple SNPs that significantly alter gene expression (Fig. 2A). 439 significant candidate SNPs impacting 76 genes across 124 eQTL were brought forward for validation.

**Figure 2.**
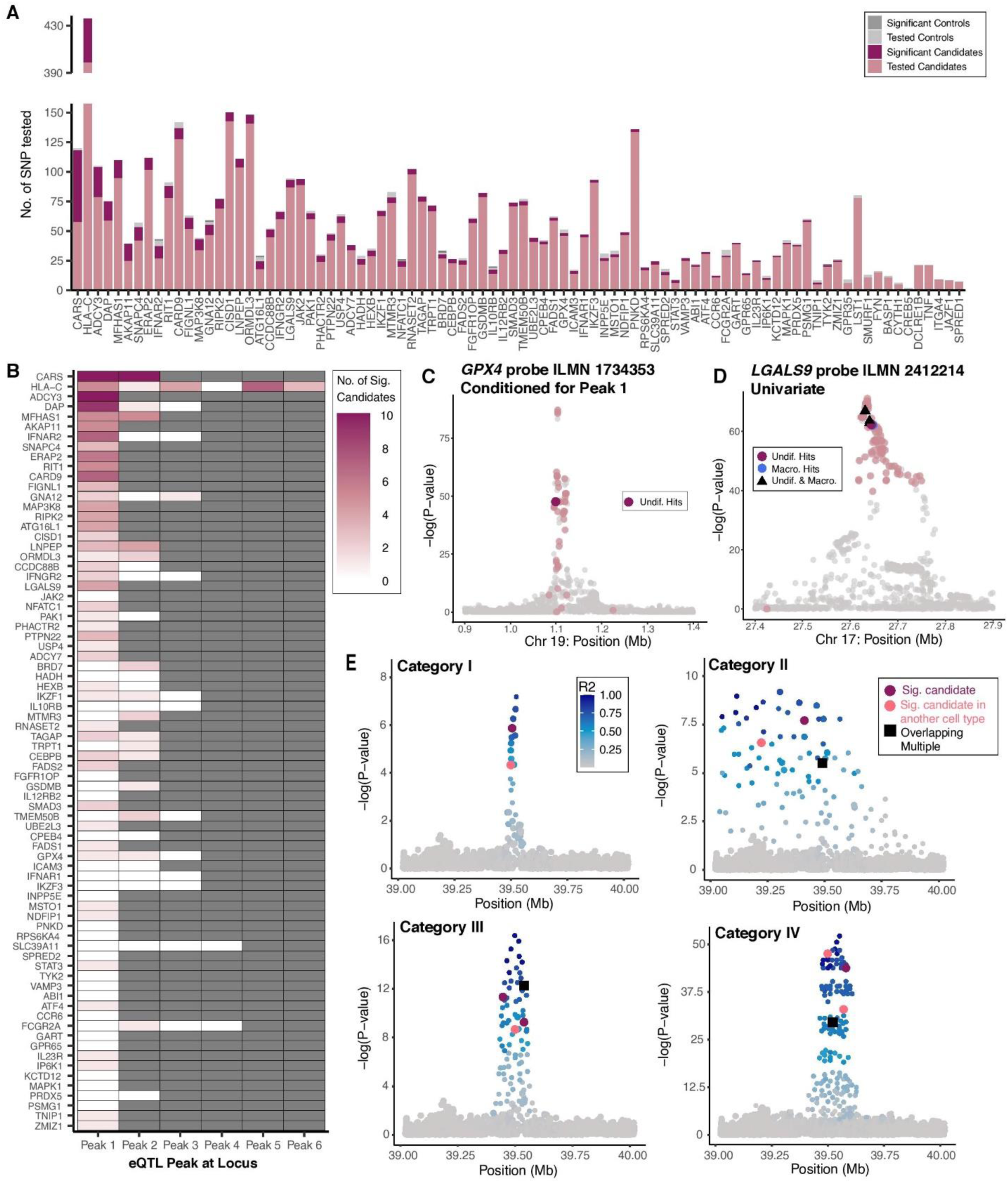
Results of Initial Screening and Validation Experiments. (A) Barplot of 87 genes tested in initial common variant experiments encompassing 4,382 gRNA across 50 transfections. A few SNPs were found in eQTL for multiple genes, so we ultimately investigated 4,814 SNP-gene regulatory relationships. Tested but nonsignificant candidates for a target gene are in salmon (n=4,272), significant candidates in maroon (n=432), nonsignificant negative controls in light grey (n=103), and significant negative controls in dark grey (n=7). (B) Heat map reveals the total number of significant candidates across all three cell types per eQTL peak following validation experiments for 76 genes brought forward. Each locus is separated into the number of peaks detected at that locus following all-but-one conditional analysis. Dark grey blocks indicate that peak number was not present at the locus. The darkest maroon indicates more than 10 significant candidates associated with that peak. (C) Custom LocusZoom plot highlighting the eQTL for GPX4 with log transformed significance on the y-axis and chromosomal position (GrCh38/hg38) on the x-axis. Significance reports the level of association with gene expression following all-but-one conditional analysis. Each point is a variant is shaded based on eCROPseq results with salmon points being all variants tested, but not significant. Maroon points indicate candidates that were validated in undifferentiated HL-60 only. (D) Custom LocusZoom for LGALS9, with same shading as in C with the addition of blue points that represent candidates that were significant in macrophage only and upwards black triangle that indicate a variant that was significant in both undifferentiated and macrophage. (E) Schematic showing four eQTL Categories and respective LD and significant candidate patterns. Darker blue shading indicates higher LD with the lead variant. Maroon points highlight significant candidates in one cell type, coral points highlight significant candidates in another cell type, and black squares highlight variants that were significant in multiple cell types.

Validation was performed in 12 transductions, each with approximately 50 previously significant candidates as well as a series of positive and negative controls (see Methods). Transcript abundance was measured with Fluent Biosciences (28) to ensure reproducibility across platforms. Guides targeting 416 out of 439 candidate SNPs were detected, of which 99 (23.8%) were nominally significant (p-value < 0.05) with a false positive rate of 7.4%. Within this group of significant candidates, only three were lead eQTL variants, for the genes *IFNAR2*, *ATF4*, and *BRD7* (Fig. 3D, Fig. 4). Significant candidates were enriched for overlap with ATACseq peaks in HL-60 cells (32) when compared to the overlap of all candidates included in our validation screening, 20.8% and 5.8% respectively (table S1). Results with alternate analytical strategies that increase power at the expense of a higher false positive rate are provided in Fig. 5.

**Figure 3.**
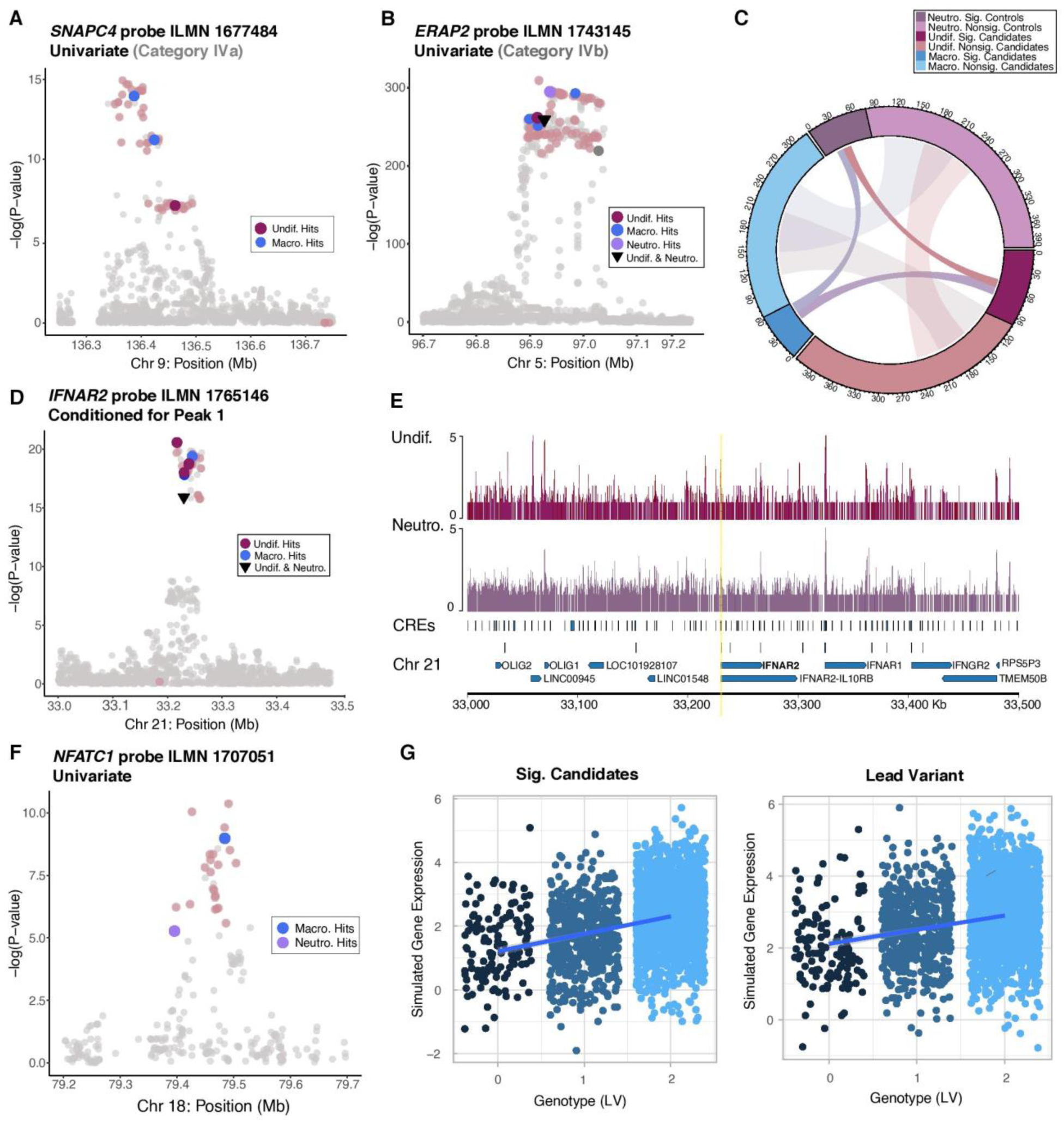
Overlapping Significant Variants Between Cell Types. (A) A) An example of Category IVa shown on a custom LocusZoom (56) plot for the eQTL of SNAPC4. Points are highlighted based on eCROPseq results with salmon points being all variants tested, but not significant. Maroon points indicate candidates that were significant in undifferentiated HL-60 only and blue in macrophage only. (B) An example of Category IVb shown on a custom LocusZoom of ERAP2, with same shading as in A. In addition, purple indicates significant candidates in neutrophil, downwards triangle indicates a variant that was significant in both undifferentiated and neutrophil, and dark grey a significant control. (C) Circos plot shading along the ring shows the number of tested candidates, but not significant (lighter color) and significant candidates (p-value < 0.05) (darker color) for undifferentiated HL-60 in maroon, macrophage in blue, and neutrophil in purple. The size of darker connections indicates the number of significant candidates detected following eCROPseq overlapping between each cell type: 18 between undifferentiated and macrophage, 14 between undifferentiated and neutrophil, and 16 between macrophage and neutrophil. Five SNPs overlap between all three cell types. The size of the larger translucent connections indicates the number of true causal candidates estimated statistically to overlap in each pairwise comparison: 121 between undifferentiated and macrophages, 102 between undifferentiated and neutrophils, and 133 between macrophages and neutrophils. (D) Primary eQTL detected for IFNAR2 with same shading as in A and B. In total we detected 7 significant candidates across the three cell types for peak 1. Specifically, one was significant in both undifferentiated HL-60 and neutrophil (rs17860115), four were specific to undifferentiated HL-60, and two were specific to macrophage. (E) Rs17860115 location highlighted in yellow revealing overlap with ATACseq peaks in undifferentiated HL-60 cells in the first track, ATACseq peaks in 120hr differentiated neutrophils in the second track, and reported cis-regulatory elements from ENCODE and NCBI in the third track. Rs17860115 overlaps with a promoter like signature reported in ENCODE and predicted silencer. The fourth track shows the position of IFNAR2 with respect to the variant. Rs17860115 falls into the 5’ UTR of the gene. (F) Primary eQTL peak for NFATC1, following eCROPseq with same shading as previously. Two candidates were found to be significant, one in macrophages (rs9748916), the other in neutrophils (rs4799052). (G) Variance explained by significant candidates (rs9748916 and rs4799052) with effect sizes of 0.5 and −0.4, together these variants explain as much variance as the lead variant (rs9962906) with effect size 0.45. All positions reported are with respect to the GrCh38/hg38 human reference genome.

**Figure 4.**
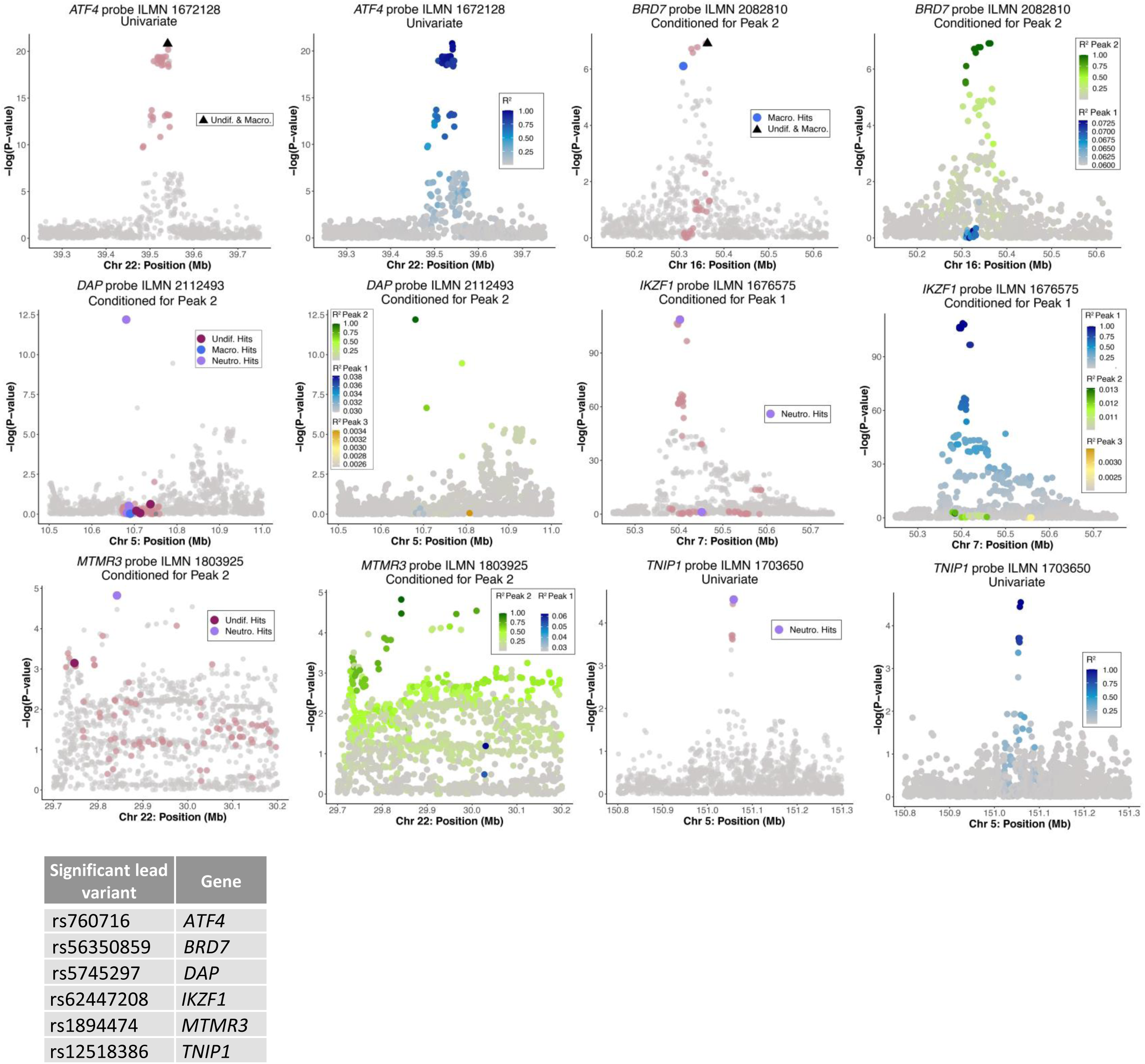
Expression QTL of Genes with Significant Lead Variants Detected in eCROPseq. Custom LocusZoom plots for genes that we detected a significant lead variant. Each gene has two plots: the left highlighting eCROPseq results and the right highlighting linkage disequilibrium (LD) patterns. On the left plot, SNPs tested are in salmon and significant candidates for undifferentiated are in maroon, macrophage in blue, and neutrophils in purple. The corresponding LD plots, on the right, highlight LD patterns for peak one in blue, with darker shading indicating higher LD with the lead variant. *BRD7*, *DAP*, *IKZF1*, and *MTMR3* had multiple peaks at the locus and whichever peak the significant lead variant was associated with is visualized (i.e. peak 1 for *IKZF1* and peak 2 for *BRD7*, *DAP*, and *MTMR3*). LD patterns for peak 2 are shown in green and yellow for peak 3. Positions are reported with respect to the hg38 reference genome. Table reports significant lead variant for the corresponding gene shown.

**Figure 5.**
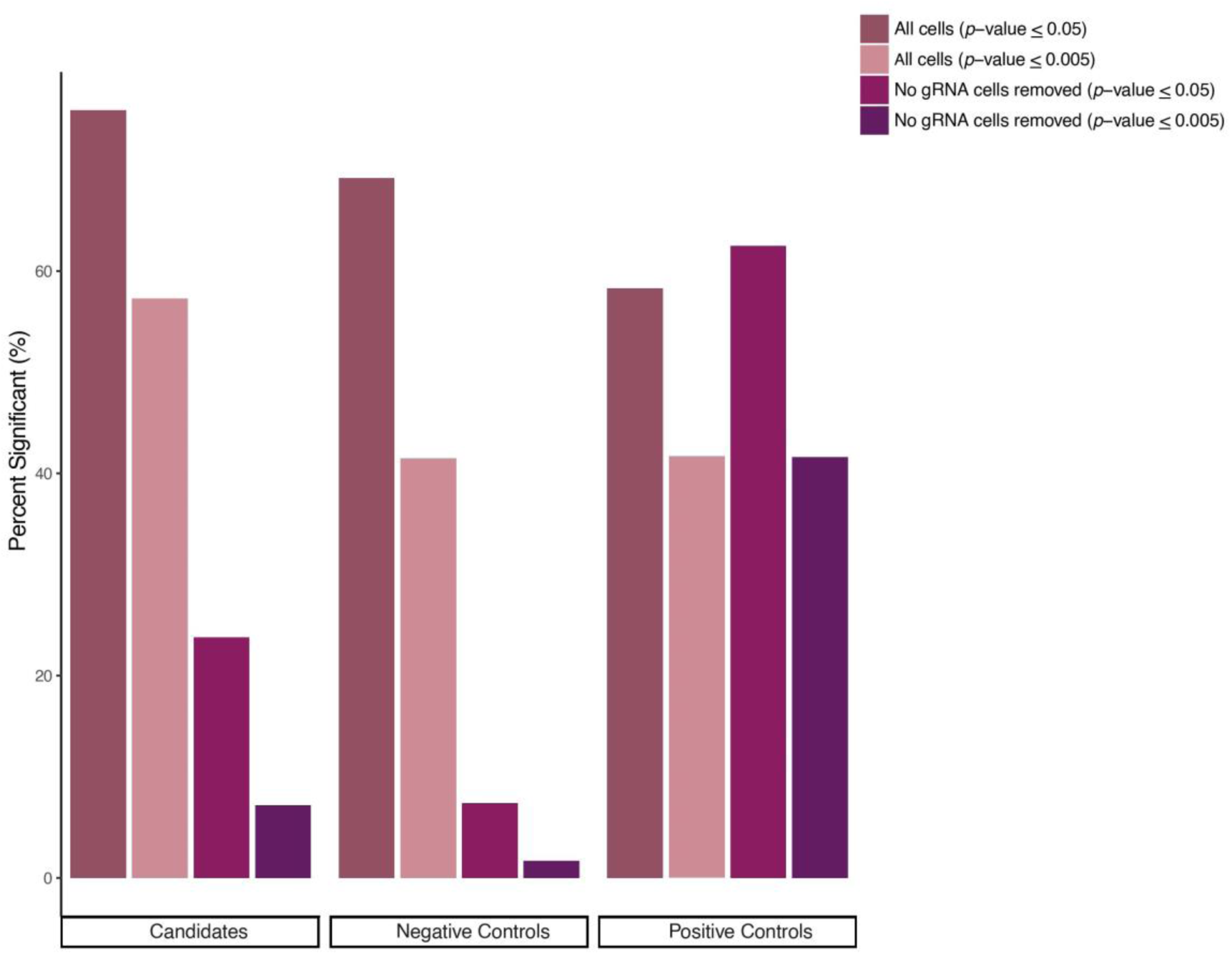
Results of Alternative Hypothesis Testing Compared to Analysis Reported. Bar plot reports proportion significant for each variant category: candidates, negative controls, and positive controls of *CISD1* and *PARK7*. Analysis with all cells is shown in brown and light pink and analysis after the removal of cells without a detectable gRNA is shown in maroon and purple. All results for rare and common variant validation are reported following analysis with the removal of cells without a detectable gRNA.

The validation set almost certainly includes a portion of true negatives due to our liberal inclusion criteria. We can estimate this portion from the data by utilizing the called positive rate (23.8%), specificity (92.6%), and sensitivity (62.5%), implying that only 30% of the variants are true positives, with a precision of 78% (see Methods). However, the positive controls are likely to have larger, more measurable effects than the candidates as these are validated SNPs known to overlap with ATACseq peaks in contrast to the lower rate of overlap observed in candidates (table S1). A better estimate with half the variants as true negatives indicated in Table 1 has a sensitivity of 40.5%, yielding a positive predictive value of 85%, a negative predictive value of 61%, and an overall accuracy of 0.67. In this case, 84 of the estimated 208 true causal variants are correctly identified and 124 are missed.

**Table 1.** Accuracy of eCROPseq Analysis in Undifferentiated Samples.

| 1:1 Ratio of True Causal Candidates to Falsely Identified Candidates | Undifferentiated: |  |  |  |
| --- | --- | --- | --- | --- |
|  |  | True Causal Candidates | Falsely Identified Candidates | Predicted Values (PPV/NPV) |
| **Specificity** 93% | **Called Positive** | 84 | 15 | 0.85 |
| **Sensitivity** 40.5% | **Called Negative** | 124 | 193 | 0.61 |
| **Sample Size** 416 | **Accuracy** 0.67 |  |  |  |

Regardless of the true proportion of negatives, the high positive predictive value in any tested scenario verifies that individual eQTL signals typically fine-map experimentally to more than one causal SNP, implying that eQTL signals are more complex than generally assumed (33). In 40.5% of all instances with a significant candidate in undifferentiated samples, we observed multiple hits spread throughout a locus and highly concentrated to a specific eQTL peak (Fig. 2B), suggesting regulation may be shaped by haplotype effects instead of individual SNP effects. For *GPX4*, we observe a signal causal candidate (rs4806970) (Fig. 2C), and for *ADCY7*, we detected two significant candidates (rs3813755 and rs34222100) (Fig. 6), all three of which have been previously associated with IBD and CD (34,35). We also identify three causal variants for *LGALS9* (Fig. 2D), two (rs3751093 and rs4794975) of which have already been associated with galectin-9 levels (36). The results are clearly driven by the impact of functional candidate variants on target gene expression, as we see limited impact on proximal and non-proximal genes (Fig. 7).

**Figure 6.**
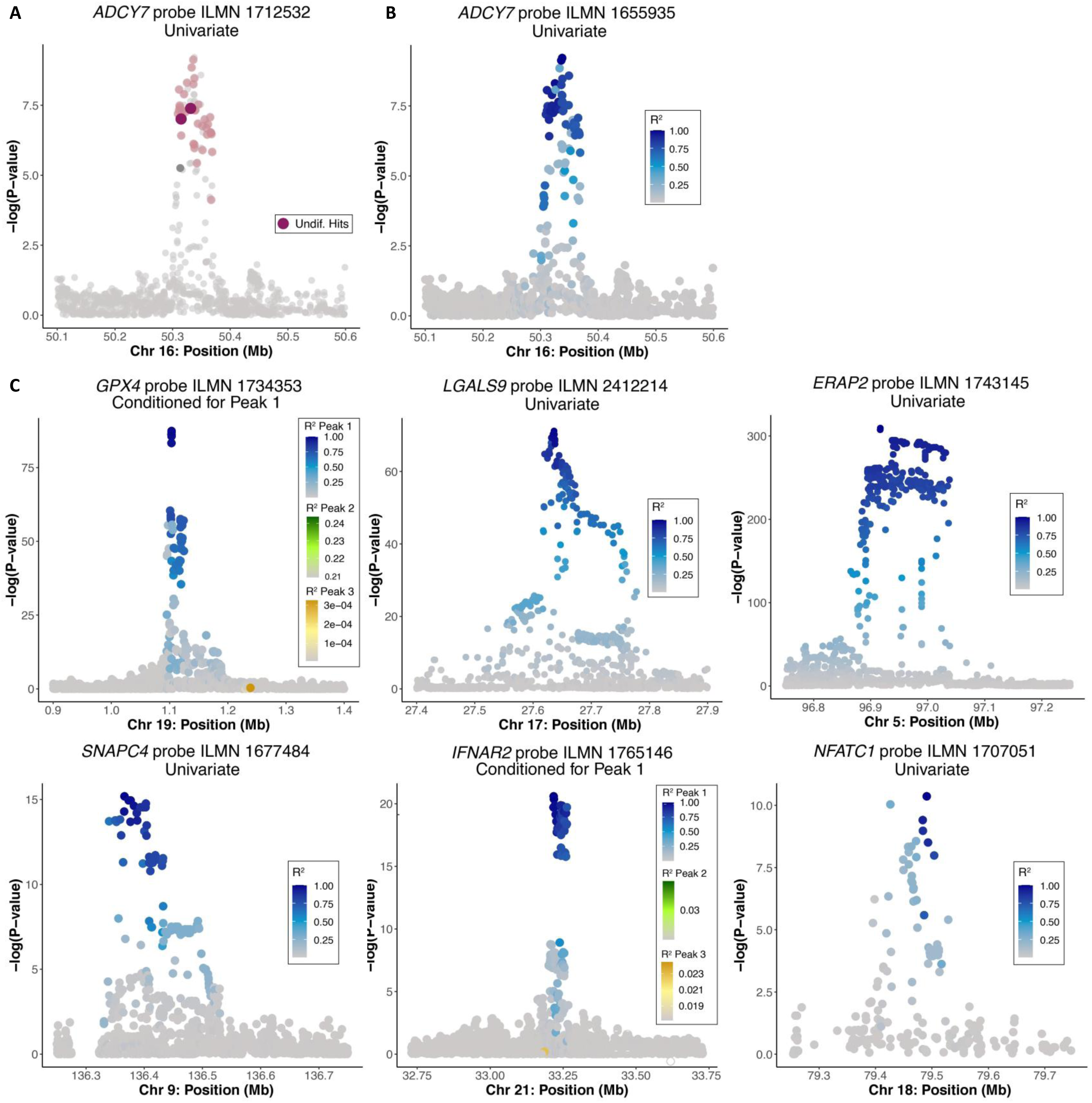
Expression QTL Highlighted by Linkage Disequilibrium. A) Custom LocusZoom of *ADCY7* highlighting eCROPseq results. SNPs tested are in salmon, significant candidates for undifferentiated are in maroon, and a significant negative control in dark grey. B) *ADCY7* plot shaded by linkage disequilibrium (LD) with darker blue shading indicating higher LD with lead variant. C) Custom LocusZoom highlights LD for all genes explicitly mentioned in text. Shading is the same as B for genes with a single eQTL peak. For genes with multiple eQTL peaks (*GPX4* and *IFNAR2*), the first peak is shown but variants associated with peaks 2 and 3 are visible on peak 1 and are shaded in green and yellow, respectively. Positions are reported with respect to the hg38 reference genome.

**Figure 7.**
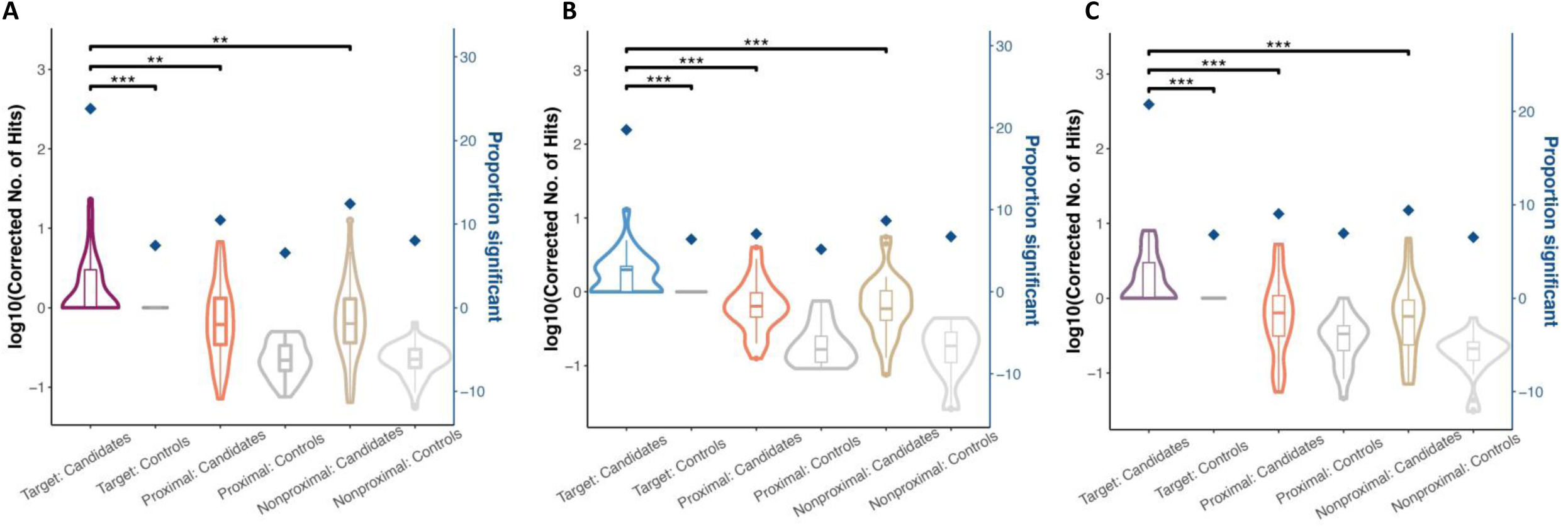
Target, Proximal, Nonproximal Analyses Across Cell Types in Common Variant Pools. Plots compare the log normalized number of significant hits after filtering for target genes with at least one significant candidate or control. A) Analysis in Undifferentiated. B) Analysis in Macrophage. C) Analysis in Neutrophil. The number of significant hits altering target gene expression is visualized in the first two violin plots. Significant candidates for undifferentiated shaded in maroon and significant controls in dark grey. Similarly, significant candidates for macrophage in blue, significant controls in dark grey and significant candidates for neutrophil in purple, significant controls in dark grey. The middle two violin plots in A,B, and C report the number of significant candidates altering proximal gene expression (orange) and significant controls altering proximal gene expression (medium grey). The last two violin plots in A,B, and C report the number of significant candidates altering nonproximal gene expression (tan) and significant controls altering proximal gene expression (light grey). The number of hits for the proximal and nonproximal genes were corrected by dividing by the total number of proximal and nonproximal genes tested, respectively. The blue diamonds indicate the percent of variants that were significant for each group, out of all those tested. Denoted comparisons between groups are following Wilcoxon rank sum test with the following ***=0.001, **=0.01, *=0.05.

### Assessing Causal Variant Specificity in Differentiated Macrophages and Neutrophils

Cell type specific eQTL effects have been reported^37–39^, but the extent of these effects is understudied. Thus, we sought to quantify the overlap between causal variants in the undifferentiated HL-60 compared to two derived cell types. The abovementioned 12 edited HL-60 validation pools were split and differentiated into macrophage and neutrophil-like cells (figs S5 and S7). We identified 64 significant SNP disruptions in macrophages and 82 in neutrophils (p-value < 0.05) (Fig. 3C). The sensitivities and specificities in each cell type are similar to that previously observed in undifferentiated HL-60 pools: 60% and 93.6% in macrophages and 50% and 93.2% in neutrophils. Combinations of significant candidates across different cell types can also explain just as much variance in target gene expression as the single lead variant of a peak^40^. For example, by simulating gene expression (see Methods), we show that significant candidates, rs9748916 and rs4799052 (Fig. 3F), explain more variance in *NFATC1* expression (*R*^2^= 6.5%) at effect sizes of 0.5 and −0.4, respectively, than the lead variant rs9962906 at an effect size of 0.45 (*R*^2^=6.1%) (Fig. 3G).

We identified a significant candidate in at least one cell type for 52 of 76 genes, including 66 eQTL peaks with clear enrichment in the primary peak (Fig. 2B). Dividing these eQTL into four categories based on size, peak height, and LD patterns, we observe trends in the location and number of causal SNPs per eQTL category (Fig. 2E). Category I (15% of cases) classifies eQTL that covers less than 200kb with few associated SNPs at an approximate max negative log p-value (NLP) between 5 12. For these peaks, we usually observe between one to two significant candidates in the middle and upper part of the peak. Category II eQTL (20% of cases) are stronger with reported eQTL signals between 5-31 and cover a much longer distance between 500-1000kb with many variants in high LD throughout. We usually observe one to four significant variants throughout the region with as many as 13 detected for *ADCY3*, suggesting unique susceptibility to perturbation. Category III eQTL (23% of cases) have more associated variants than category I and are evenly dispersed throughout the peak with a gradual decline of LD over a region of 200kb and signal NLP>10 for the majority of eQTL in this category. We often detect several hits towards the top or middle of the peak as in eQTL for *ADCY7* (Fig. 6A), with an average of three per peak, as well as numerous instances of overlapping significant candidates between cell types. For example, two *LGALS9* candidates (rs3751093 and rs4795830) were also found significant in macrophage samples (Fig. 2D).

Category IV are the most significant signals with the majority of peaks having a max NLP>20. These peaks generally cover a distance between 200-300kb and have breaks in the eQTL creating individual tiers of SNPs at distinct significance and LD levels (Fig. 6C). We subdivide this category into category IVa (21% of cases) and category IVb (21% of cases) based on whether tiers harbor significant candidates from multiple cell types or not. Category IVa tiers are driven by distinct cell types as shown in *SNAPC4* (Fig. 3A) whereas category IVb has multiple cell types and overlapping significant candidates as observed in ERAP2 (Fig. 3B). Additionally, the previously mentioned significant lead variants for *ATF4* and *BRD7* were also found to be significant in macrophages. Lead variants for four genes (*DAP*, *IKZF1*, *MTMR3*, *TNIP1*) were significant in neutrophils (Fig. 4), half of which were found associated with the secondary eQTL peak, suggesting that multiple eQTL peaks at a locus in whole blood may each be driven by a specific cell type instead of representing a shared signal between multiple cell types as may be the case for the primary peak.

Contrasting across cell types, we observe 18 overlapping significant candidates from 9 genes between undifferentiated HL-60 and macrophages; 14 from 10 genes between undifferentiated HL 60 and neutrophils; and 16 from 9 genes between macrophages and neutrophils (Fig. 3C).

Significant candidates in two or more cell types are also enriched in reported regions of open chromatin (table S1). One example, rs17860115, altered *IFNAR2* expression in both undifferentiated HL-60 cells and neutrophils (Fig. 3D), overlaps a region with a promoter like signal^41^, and a predicted silencer^42^ in human lymphoblastoid cells (Fig. 3E), which can explain the decrease in expression observed when targeted. In addition, we identify two macrophage-specific and four undifferentiated HL-60 specific causal candidates, one of which is the lead variant for the primary peak (rs56079299). Five out of these seven variants have also been implicated in severe COVID-19^43^ highlighting the clinical applicability of expression CROPseq.

Of all overlapping causal candidates, only 5 overlapped across all three cell types and altered expression for one of four genes: *HLA-C*, *LNPEP*, *MFHAS1*, and *PAK1*. Following our approach above to estimate sensitivities, we then predict the proportion of overlapping true causal candidates in each pairwise contrast (see Methods, tables S2-S4). Between undifferentiated HL-60 and macrophage as well as macrophage and neutrophils, the proportion of overlap is inferred to be 75 and 87%, but this drops to 54% between undifferentiated and neutrophil (Table 2). This suggests that the shared regulatory networks between immune cells is likely much greater than previously reported, and by extending CRISPR screenings to more cell types, evaluation of the nature of these related landscapes will unfold.

**Table 2.** Estimated percent overlap of true causal candidates per cell type comparison.

| Comparison | Undifferentiated (Cell Type 1) and Macrophage (Cell Type 2) | Undifferentiated (Cell Type 1) and Neutrophil (Cell Type 2) | Macrophage (Cell Type 1) and Neutrophil (Cell Type 2) |
| --- | --- | --- | --- |
| No. of significant candidates overlapping | 18 | 14 | 16 |
| Sensitivity of Cell Type 1 | 0.45 | 0.41 | 0.34 |
| Sensitivity of Cell Type 2 | 0.33 | 0.335 | 0.355 |
| No. of true causal candidates | 161 | 190 | 152 |
| Estimated no. of overlapping true causal candidates | 121 | 102 | 133 |
| Estimated percent overlap of true causal candidates | 75% | 54% | 87% |

## DISCUSSION

Here, we provide a proof-of-principle that eCROPseq is an efficient way to test the functionality of rare variants, which may be particularly useful for rapid clinical testing. We also show the effectiveness of eCROPseq in fine mapping common variants in eQTL, and establish that by screening across credible sets and surrounding SNPs in high LD, individual eQTL peaks often map to multiple candidate sites rather than reducing to a single causal variant. Across 66 eQTL with at least one significant candidate following our common variant screening, we observe an average of 2.3 causal candidates across all three cell types tested: undifferentiated HL-60 cells, macrophages, and neutrophils. The most direct interpretation is that multiple SNPs acting together are responsible for the regulatory influence, within which two or more SNPs are functional. While it is well established that the expression of many genes is influenced by multiple local (cis-acting) eQTL^4,8^ in low LD, the results reported here suggest an added layer of complexity with each independent eQTL often explained by a complex haplotype effect. Further, it appears that some functional variants within each credible set may be cell-type specific, with others combining to generate the statistical signal in an individual cell type.

An alternative interpretation is that the CROPseq assay, which is based on inducing small indels, over-estimates the likelihood that a transition or transversion at the same site influences gene expression. Our results for both the common variant and rare variant datasets may be due to perturbation altering transcription locally, without directly informing what proportion of single nucleotide changes have the same effect. Comprehensive base-editing^44^ of multiple causal variants singly and in combination will be required to distinguish between these two interpretations. However, recent base-editing evidence suggests that observed effects are attributable to haplotype effects. Specifically, one base-editing screen^45^ confirmed that multiple putative causal variants in a 100bp promoter element of *HBG1*/*2* disrupt expression. Similarly, we conducted a pilot adenine base-editing study of our positive hits at 3 *DAP* and 4 *CARS* SNPs (Fig. 8), validating significant reduction of expression relative to negative controls at all sites, implying that substitutions as well as microdeletions have measurable effects on gene expression. More experiments of this type are underway to quantify the general functionality of nucleotide substitutions.

**Figure 8.**
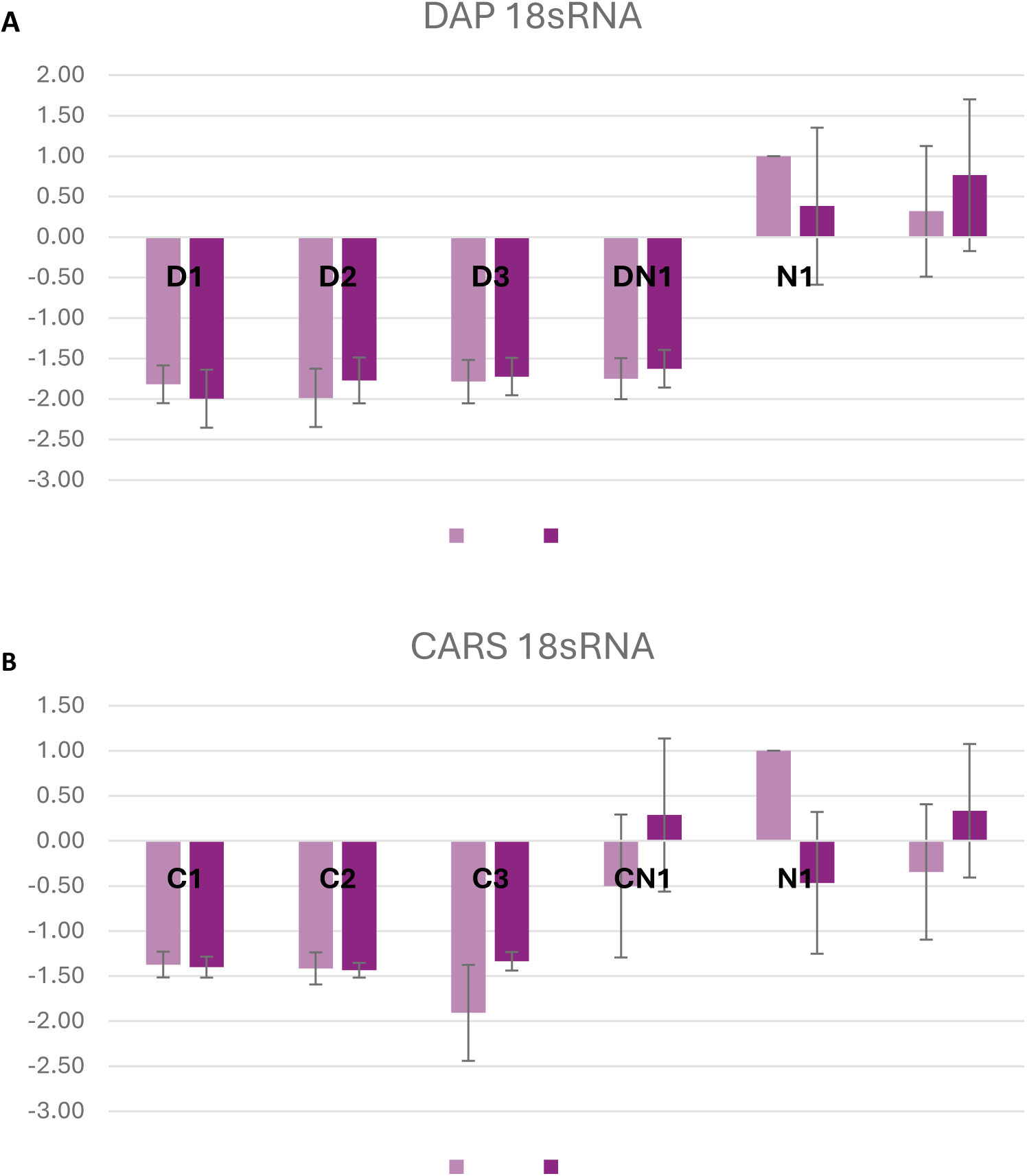
Decrease of DAP and CARS Expression Following Base Editing of Candidates. qPCR bar plots show relative expression of DAP (A) and CARS (B) in base-edited cells (D1-D3, DN1, C1-C3, CN1) and non-targeting negative controls (N1 and N2). D1 (rs296479) and D2 (rs2918392) are significant candidates validated from eCROPseq validation. D3 (rs10866479) was significant in initial experiments and in analysis that included all cells, but not in the analysis with removal of cells without a detectable gRNA. DN1 (rs12654966) was not significant in eCROPseq. C1-C3 (rs10488673, rs7929049, and rs440130) are validated significant candidates while CN1 (rs756693) was not significant in eCROPseq. Two different multiplicity of infections (MOIs), low (light pink) and high (dark pink), of lentivirus transduced base editing were also compared. Base-editing efficiency was confirmed through ICE analysis. Each sample contains three technical replicates. qPCR Ct values were normalized against a housekeeping gene, 18sRNA and the relative expressions were calculated by the ΔΔCt method and scaled to N1-low as 1 of each gene. Direction of effects for candidates match the direction observed in the single-cell data.

If our results are confirmed, the implication that haplotype effects commonly explain eQTL would be considerable. Our characterization of four classes of eQTL underlines the complexity observed even at single peaks, which often show dozens of variants in tiers of high LD with variable degrees of overlap. Similarly, GWAS signals display diverse patterns with tiered LD and overlapping signals, but there is little theory explaining why. Selection on gene expression appears to be weak and mostly stabilizing (46), to which we would add a high likelihood of softness^19,20,47^, namely multiple variants co-evolving rather than a single causal site with a large effect size. While it is not clear to what extent eQTL and GWAS trait associations colocalize^48^, dispersion of haplotype effects over, in some cases, hundreds of kilobases further obscures the relationship. The prospect that regulatory signals are due to multiple variants in credible sets call for more research on the implications for the architecture of QTL. Notably as well, variation across ancestry groups in the number of SNPs contributing and their exact LD patterns may also explain heterogeneity of allelic effect sizes, and of the colocalization of eQTL and GWAS signals. This has implications with respect to the predicted efficacy of somatic or germline gene editing, as well as for understanding the low transferability of polygenic scores^21,22^. Investigation of haplotype effects in different primary cells from individuals with diverse genetic backgrounds could elucidate variability of regulatory effects across different ancestry groups.

Major limitations of this study are that the eCROPseq strategy is only powered to detect up to half of the functional effects, that we have not accounted for likely variability in cutting efficiency at each site by incorporating multiomic ATAC and RNA sequencing on the same cells^49^, and that large scale base editing will be needed to affirm that deletion effect generally mirror nucleotide substitution effects. We focused on myeloid cells in this study given possibilities to examine complex differentiation during the innate response, but comparison of regulatory overlap with adaptive immune cells will be of interest in future experiments. Nevertheless, at a minimum, our results imply that variant-to-function studies^5^ ought not concentrate solely on lead variants. Rather, the assumption that single causal variants explain eQTL and by extension GWAS signals needs to be rigorously evaluated experimentally.

## Materials and Methods

### Rare Variant Selection and gRNA Design

To select variants for analyses, we first selected target genes that were expressed in our Cas9 integrated HL-60 cell line (fig. S2). We then selected candidate rare variants (MAF < 1%) that had a Watershed^27^ posterior probability greater than or equal to 0.9. For each gene, we also selected four controls predicted to be non-functional with a Watershed posterior probability less than or equal to 0.01. We further filtered our gene set to include those with at least five rare variants predicted to be functional and designed two unique 19 base pair gRNA. Both targeting gRNA were present in the scRNAseq data for almost all of the rare candidates. Each control was targeted by one unique gRNA. Rare candidate and control gRNA were designed with the following criteria: low repetitiveness within the range of 20-80% GC content, nearby PAM sequence (NGG), a cut site 10 or less base pairs from the reported variant position. The majority of gRNA had minimal off targets predicted by COSMID (50) with searching based on GRCh38/hg38 reference genome and criteria of no indels or 1 base pair deletions or insertions. This resulted in 465 rare candidates and 327 controls across 89 genes, averaging 5.2 rare candidates and 3.7 controls for each gene considered. A comprehensive list of gRNA and corresponding target genes used in the analysis is reported in table S6.

### IBD Gene Set Selection, eQTL Isolation, and Common Variant Selection

Target gene selection and eQTL isolation through all-but-one conditional analysis was detailed in our previous work^30^. Briefly, 276 genes associated with inflammatory bowel disease (IBD) were selected based on their implication in one or more genome wide association studies (GWAS) for IBD and reported on Immunobase and OpenTargets (https://genetics.opentargets.org/immunobase accessed January 2019 and https://www.opentargets.org accessed November 2021). Cis-eQTL for these genes were searched within a 1Mb region surrounding the TSS of the gene (+/- 500kb) using the Consortium for the Architecture of Gene Expression peripheral blood microarray data from 2,138 individuals of European descent^51^ and isolated using all-but-one (AbO) conditional analysis. 165 genes had at least one significant eQTL after AbO conditional analysis of which 87 were expressed in our HL-60 cell line and tested during our eCROPseq common variant assay. All eQTL profiles can be browsed on our Shiny application at https://eqtlhub-gt.shinyapps.io/shiny/

For each of these genes, we manually selected candidate SNPs by choosing credible sets and surrounding SNPs that were in high LD with the lead variant, strongly significant in the peak, or overlapped reported ATACseq peaks in HL-60, Jurkat T-cells, or Monocytes (GSM2083754^52^, GSM4005276^53^, and GSM2679893^54^). GSM2083754 was transformed to GRCh37/hg19 reference genome using UCSC’s liftover^55^. We also selected negative controls for each gene which were found outside all eQTL signals present at the locus. Variants were then checked for targetability with the same criteria mentioned in Rare Variant Selection and gRNA Design (low repetitiveness, nearby PAM sequence and cut site). The majority of gRNA had minimal predicted off target effects from COSMID^50^ based on searching with no indels, deletions or insertions in the GRCh37/hg19 reference genome. All candidate and control SNPs were targeted with one unique gRNA per variant.

Unique guide RNAs for each candidate and control was cloned into oligonucleotides and split across 50 lentiviral transductions (see Oligonucleotide Synthesis and Lentiviral Transduction). Cells were prepared for scRNAseq using 10x genomics 3’ v3.1(CG000204, 10x Genomics) and processed following the workflow mentioned below. We profiled an average of 15,239 HL-60 cells after transduction, allowing adequate counts to characterize gene expression changes. Ultimately, 4,382 variants were targeted across 87 IBD associated genes. A handful of variants were included in eQTL for more than one gene, so a total of 4,814 regulatory relationships were tested. All SNPs and corresponding gRNA are listed in table S7.

Of the 4,382 variants tested, 439 were significant (p-value < 0.005) (see Hypothesis Testing in Seurat) across 76 genes and were brought forward to validation. Tested and significant candidates for eQTL were transformed to the GRCh38/hg38 reference genome via liftover^55^ and highlighted on custom LocusZoom^56^ plots in RStudio.

### Controls

Positive and negative control gRNA were included in every plasmid library. In our initial common variant screening, every pool contained a minimum of four controls including one negative control that had no perfect target in the human genome, another negative control that targeted a non-SNP region, and two positive controls impacting cell survival by disrupting *RUNX1* or *TUBB*. Rare variant pools included 9 controls consisting of those previously listed as well as three more positive controls targeting *RUNX1* or *TUBB*, and two previously eCROPseq validated common variants impacting *CISD1* and *PARK7* (rs2251039 and rs35675666)^25^. Common variant validation pools had a total of 8 controls including the non-SNP negative control, two non-targeting controls, three positive controls, as well as the CISD1 and PARK7 controls. A comprehensive list of the controls used throughout the eCROPseq assays, and the corresponding gRNA sequences, are listed in table S5.

Negative controls and positive controls that impact cell survival were used to verify sufficient editing from Cas9 by comparing the proportion of gRNA in a plasmid library to that of the scRNAseq library (fig. S1). The proportion of a gRNA in a respective plasmid library was calculated by the number of reads mapping to a specific gRNA divided by the total number of reads mapping to all other gRNA in the library. The proportion of a gRNA in a quality controlled, normalized, and scaled scRNAseq library was calculated by the number of cells containing a specific gRNA divided by the total number of cells containing all other gRNA in the pool. Positive controls are expected to show fewer counts in the scRNAseq library since they target genes necessary for survival (*RUNX1* and *TUBB*), thus cells will die if editing occurred, which will be reflected in the data as a decreased proportion in the scRNAseq library when compared to the proportion in the plasmid library. This was observed in the vast majority of pools. As expected, negative controls showed minimal differences in the proportion between the two libraries because they targeted nonSNP regions or had no match in the human genome, thus no editing occurred in regulatory regions or at all.

### Guide RNA Library Design, Cloning and Validation

We added a G to the first base of the gRNA sequence to increase editing efficiency^57^. Afterward, 18 bp and 35 bp of homology to the hU6 promoter and gRNA backbone were added to each designed gRNA (5’-TGGAAAGGACGAAACACCG - gRNA - GTTTTAGAGCTAGAAATAGCAAGTTAAAATAAGGC-3’) for downstream Gibson Assembly. These gRNA libraries were then synthesized by Twist Bioscience ^58^ and were suspended (100nM) in TE buffer (Qiagen) for use. The CROPseq-Guide-Puro plasmid^24^ (Addgene, 86708) was digested by restriction enzyme, Esp3I (NEB, R0734L) and gel-purified (Qiagen, 28706) to extract the backbone fragment (8,333 bp).

Gibson assembly reactions were performed to insert gRNA (200 fmoles) into the plasmid (11 fmoles) using NEBuilder HiFi DNA Assembly Master Mix protocol (NEB, E2621S). Reaction products were transferred onto a membrane filter (Merck, VMWP04700), floating on HPLC-quality water for 30 min to de-salt, and then electroporated into Endura electrocompetent cells (Lucigen, 60242-1). Electroporated cells were coated on LB-agar petri dishes (Amp) and incubated overnight (16-18 hours) at 37°C for Maxi-prep (ZymoPURE II Plasmid Maxiprep Kits, D4203). The gRNA distribution in each plasmid library was validated by next-generation sequencing (NGS) on an Illumina MiSeq. In over 60% of pools, we detected just 1 gRNA in the majority of cells (fig. S8).

### Lentivirus Library Construction, Transduction, and Puromycin Selection

Lentivirus was produced using the 2nd generation lentivirus system, with the constructed gRNA plasmid (CROPseq-Guide-Puro), an envelope plasmid, pMD2.g (Addgene, 12259) and a packaging plasmid, psMAX2 (Addgene, 12260). Following Addgene’s standard lentivirus production protocol, the mixture of the 3 transfection plasmids was prepared (DNA: PEI ratio = 1:3) in Opti-MEM I reduced serum medium (Gibco, 31985062) and added to Lenti-X 293 T cells (Takara, 632180) for virus production, cultured in DMEM medium (Sigma, D5796-6X500ML) with 10% fetal bovine serum (FBS, Corning, 35-010-CV) and 4 mM L-glutamine (Gibco, 25030081). pLenti-CMV-GFP-Puro (Addgene, 17448) was also produced as the reporter in each transfection. Collected lentivirus was concentrated using Lenti-X Concentrator (Takara, 631232) and dissolved in 500 μl 1x DPBS (Gibco, 14190-094). Lentivirus titers were measured by qRT-PCR using Lenti-X qRT-PCR Titration Kit (Takara, 631235).

An HL-60/S4-Cas9 cell line, which can produce Cas9 protein, was generated as described in our previous study^25^. The Cas9 gene integration was validated by ddPCR and functionality was validated by nucleofection with a gRNA, R-66S targeting the *HBB* gene, followed by Sanger sequencing and Synthego ICE analysis (Synthego Performance Analysis version 3.0). We used spinfection to deliver lentivirus to the cell line according to a CRISPR-Cas9 screening protocol58. In brief, cells were seeded in a 24-well plate at a density of 1 x 10^6^ /ml with 8 µg/ml of polybrene (EMD Millipore, TR-1003-G) in IMDM medium (Gibco, 12-440-053) with 20% FBS and 1% P/S (Gibco, 15 140-122). Lentivirus, including the gRNA library and pLenti-GFP positive control, was separately added to cell culture with an estimated 0.2-0.3 multiplicity of infection (MOI) to ensure single lentivirus transduction of each cell with decent efficiency. Then, the cells were centrifuged at 1,200×g for 1.5 hours at 33°C and returned to a cell incubator at 37°C.

Twenty-four hours after spinfection, cells were replated at a density of 5 x 10^5^ /ml with 0.4 µg/ml puromycin (Invivogen, ant-pr-1). The puromycin drug selection lasted for 8 days until there was no obvious change in cell viability. Every 48 hours, cell density was measured, and the medium with puromycin was replenished. The cell density was maintained under 1-1.5 x 10^6^ /ml. MOI was calculated based on cell density measurement through puro selection. After puromycin selection, cells were cultured in regular medium for at least 10 days for global expression recovery, before single cell RNA sequencing.

### Single Cell RNASeq Library Preparation and Sequencing

After allowing 10 days for global expression recovery, cells were shipped overnight from Rice University (Houston, TX) to Georgia Tech (Atlanta, GA) in a temperature-controlled container to undergo scRNAseq library preparation using either 10x Genomics Chromium Single Cell 3’ v3.1 Chemistry Dual Index (CG000204, 10x Genomics), Fluent Biosciences T100 v4.0PLUS (FB0003657, Fluent Biosciences), or Fluent Biosciences T20 v4.0PLUS (FB0005002, Fluent Biosciences) scRNAseq library preparation, depending on the experiment. 50 transfections in initial common variant experiments were processed across the 10x Genomics platform. Rare variant pools (n=2) were processed with Fluent Biosciences T100. Common variant validation pools consisting of 12 undifferentiated HL-60, 12 macrophage, and 12 neutrophil pools were processed with the Fluent Biosciences T20. Following library preparation, samples were sequenced using the NovaSeq-6000 S4 Reagent Kit v1.5 (200 cycles) (20028313, Illumina) for 100bp pair-end sequencing conducted by the Molecular Evolution Sequencing Core at Georgia Tech. Macrophage results were only reported for pools 1-9 and 11 since macrophage pools 10 and 12 had too few cells detected following sequencing and thus were excluded from all statistical analyses.

### Sequence Alignment

After sequencing, BCL files from samples processed with 10x Genomics kits were converted to fastq files using Cell Ranger’s mkfastq^31^. BCL files from samples processed with Fluent Biosciences kits were converted to fastq files using Illumina’s bcl2fastq2 (RRID:SCR_015058). Rare variant pools were processed with Fluent Biosciences T100 kits, so scRNAseq libraries were split across multiple sequencing libraries to increase sequence diversity as suggested in the protocol (FB0003657, Fluent Biosciences). Rare variant fastq files from the same scRNAseq sample were processed together as one sample downstream.

All fastq files were then aligned to a supplemented GrCh38 human reference genome (version refdata-gex-GRCh38-2020-A, 10x Genomics), which was supplemented with artificial chromosomes for each gRNA in a library. Following Datlinger et al. ^24^, artificial chromosomes included one of the above-mentioned oligonucleotides surrounded by more of the hu6 promoter (totaling 250bp) and downstream backbone (totaling 260bp) until the poly-A tail. Reference genomes for samples processed with 10x Genomics kits were supplemented using Cell Ranger^31^ mkref (version 6.02). Fastq files for 10x Genomics samples were then aligned to the supplemented reference genome using Cell Ranger’s count and were processed in R Studio (version 4.1.2) using Seurat (version 4.4.0)^61^.

Reference genomes for samples processed with Fluent Biosciences kits were supplemented via STAR^60^ (version 2.7.11a) with commands --runMode genomeGenerate with additional commands of --sjdbOverhang 69 and --genomeSAsparseD 3. Sensitivity levels between 1-5 were manually selected for each sample by visually inspecting barcode rank plots and selecting sensitivity where barcodes representing cells end in the middle of the major drop-off of the barcode rank plot, thus eliminating unlikely cells. Filtered matrixes for the respective sensitivity threshold were processed in R Studio (version 4.1.2) using Seurat (version 4.4.0)^61^.

### Hypothesis Testing in Seurat

Filtered matrixes were read into R Studio using Seurat (version 4.4.0) and features detected in fewer than 6 cells and cells with less than 200 features were removed. In initial common variant experiments, we visually inspected each sample to determine proper QC cutoffs for the range of features and percent mitochondrial counts. Most pools retained cells that had 2,400 7,500 features and less than 25% mapping to mitochondrial reads and were split across two 10x reactions. Following QC of each individual sample, samples from the same pool were then merged together and normalized.

To improve consistency in pre-processing of rare variant and common variant validation pools, two median absolute deviations (MADS) surrounding the median of a sample’s feature count threshold and percent of reads mapping to mitochondrial content were selected as QC thresholds. Based on two MADS, the average QC threshold across rare samples was a minimum of 1,923 features, a maximum of 5,339 features, and a maximum of 6.6% of reads mapping to mitochondrial content.

The average for undifferentiated samples was a minimum of 2,422 features, a maximum of 7,482 features, and a maximum of 7.5% of reads mapping to mitochondrial counts. For macrophage, this was 1,918 minimum features, 8,108 maximum features, and a maximum of 9.8% of reads mapping to mitochondrial counts. For neutrophil, this was a minimum of 2,007 features, a maximum of 10,560 features, and a maximum of 9.8% of reads mapping to mitochondrial counts. All samples were then normalized and scaled to standard normal distribution using ScaleData(). Normalization consisted of dividing each feature count by total feature counts in a cell, log transforming, and multiplying by 10,000. After scaling, additional QC included verifying that the majority of cells received one gRNA (fig. S8) and clustered based on cell cycle phase and not based on perturbation (fig. S9).

Hypothesis testing in the initial common variant screening was conducted by comparing target gene expression between cells that received a gRNA (gRNA+) to those that did not (gRNA-) for an average of 15,222 cells per transduction. Student’s t-test rendered the most power in contrast to more advanced models in pilot experiments (25). We also observed similar power to the two sample Kolmogorov-Smirnov (KS) test, comparing the distribution in target gene expression in both groups.

Thus, for each gRNA+ and gRNA-comparison, we evaluated significance through the Student’s t-test and two sample KS test. A variant was deemed significant when the resulting p value was less than or equal to 0.005 with a minimum of five cells. We also removed cells not expressing the target gene and repeated these tests, ultimately resulting in four hypothesis tests per gRNA.

Hypothesis testing in rare variant pools was also conducted by first removing all cells without a detectable gRNA as they likely harbor a gRNA impacting a target gene of interest. This resulted in 33,302 and 43,129 cells for each pool. By decreasing cell counts in this way, we lost power (Fig. 5) so significant results of rare variant analysis are with respect to a 0.05 critical value in a minimum of five cells. Additionally, due to a candidates in the rare variant screening being targeted by two unique gRNA, we performed a series of hypothesis testing comparing target gene expression between 1) cells that received gRNA 1 (gRNA 1+) to all those that did not, 2) cells that received gRNA 2 (gRNA 2+) to all those that did not, 3) gRNA 1+ cells compared to cells that had neither gRNA 1 or 2, 4) gRNA 2+ compared to cells that had neither gRNA 1 or 2, 5) and cells that had either gRNA 1 or 2 to cells that had neither gRNA 1 or 2. We performed hypothesis testing this way for both a Student’s t-test and KS, and then repeated all tests after removing cells that did not express the target gene. All significant variants are listed in table S8.

In validation experiments, undifferentiated cells without a detectable gRNA were removed resulting in an average of 17,925 cells per pool. In macrophage and neutrophil pools, we also removed cells without a detectable gRNA as well as cells that were potentially undifferentiated. We selected high confidence differentiated cells by filtering for cells with greater than or equal to median expression or for one of two marker genes in macrophages: *CD14* and *CD68* and three marker genes in neutrophils: *CSF3R*, *MMP9*, *S100A9*. This yielded an average of 5,530 cells per macrophage pool and 4,958 cells per neutrophil pool. Marker genes were first chosen from literature and then based on those detected in our differentiated cells. UMAPs highlighting these markers in all cells that passed QC are visualized in fig. S7 which are contrasted to UMAPs in undifferentiated HL-60 pools above, revealing differential expression of marker genes. Then gRNA+ cells were compared to gRNA-cells with gRNA not targeting the same gene as these pools were enriched for previously significant gRNA target the same gene of interest, thus removing them from hypothesis testing was essential to capture the true unedited target gene distribution. This was done for both a Student’s t test and KS test. Cells not expressing the target gene were then removed and hypothesis tests were repeated in the subset of cells. Significant SNPs had a p-value less than 0.05 with a minimum of five cells in any of the 4 hypothesis tests and are listed in table S9.

Cells from undifferentiated pools 1 and 2 were kept alive and were processed as negative controls alongside macrophage and neutrophil pools. This was done to evaluate consistency after a longer culturing period and variability in scRNAseq library preparation. 40-43% of significant candidates in the control pools were found in the original pools, reflecting our estimated sensitivities (table S10).

### Assessing Background Effects

We assessed background effects of our eCROPseq assay by comparing the number of variants observed to significantly alter target gene expression to the corrected number of variants observed to significantly alter proximal and nonproximal gene expression. Proximal genes are defined as any genes in approximately a 1Mb region surrounding the TSS of the target gene (+/- 500kb) and expressed in our HL-60 cell line. Genes in this region were obtained from the UCSC Genome Browser^55^, resulting in an average of 15.5 proximal genes to a given target gene in rare variant analysis, 9.7 in undifferentiated HL-60, 8.7 in macrophage, and 9.4 in neutrophil. As for non proximal genes, a list of genes expressed in HL-60 was obtained from Harmonizome^61^ and 250 were randomly selected for rare variant screening and another 250 were randomly selected for common variant validation. Following the same statistical tests in Hypothesis testing in Seurat, we tested every gRNA for impact on those 250 genes that were adequate expressed in samples (an average of 150.5 nonproximal gene in rare variant pools, 183 in undifferentiated HL-60 pools, 161 in macrophage pools, and 167 in neutrophil pools).

The number of significant candidates for proximal genes were corrected by dividing the total number of significant candidates by the total number of proximal genes tested and by dividing the total number of significant controls by the total number of proximal genes tested. The same was done for nonproximal hits by dividing by total nonproximal genes tested. Results in Fig. 1C and Fig. 7 report results comparing target genes with at least one significant candidate.

### Neutrophil and Macrophage Differentiation for edited HL-60

The HL-60 cell line, as a human myeloid derivative, can be differentiated into neutrophils or macrophages. We also tested the 12 validation variant pools’ significance in HL-60/S4-Cas9-derived neutrophils and macrophages, respectively. The recovered cells in each pool were replated into two plates, 1 x 106 cells each. 10 µM all-trans retinoic acid (ATRA, Thermo Scientific, 207341000) was used for neutrophil differentiation, and 100 nM 1α,25-Dihydroxyvitamin D3 (Vitamin D3, Sigma Aldrich, D1530-10UG) was used for macrophage differentiation. The medium and drugs were replenished every 48 h and cell density was maintained under 1 x 106/ml. After 72 hours of differentiation, cells were stained to measure their differentiation efficiency via the BD FACSMelody cell sorter. Neutrophils are CD11b+ and undifferentiated HL-60 cells are CD11b-; macrophages are CD71-/CD14+ and undifferentiated HL-60 cells are CD71+/CD14-. Relative isotypes for each antibody were also included in cell staining. Afterward, differentiated neutrophils and macrophages were harvested for scRNAseq, and their identities were confirmed by expression of relevant marker genes.

### Overlap with regions of Open Chromatin

We also assessed the rate at which significant candidates identified in undifferentiated, macrophages, and neutrophils overlapped any reported ATACseq peaks in each respective cell type. Specifically, the number of variants that were in regions of open chromatin were detected by determining which variants had reported positions that overlap with any indicated peak in the same position. This was done for three replicates of Undifferentiated HL-60 cells (GSM2083754, GSM2083755, GSM2083756)^52^, and if a variant overlapped in at least one, it was denoted as being in a region of open chromatin. Each variant in macrophages was assessed for overlap with ATACseq peaks in HL-60 cells that were differentiated to macrophages over the course of 120 hours and were publicly available. If a variant overlapped in the region of a peak in one of the three replicates (GSM2083778, GSM2083779, GSM2083780)^52^, then it was denoted as being in a region of open chromatin. Similarly for neutrophils, HL-60 cells were differentiated to neutrophils over the course of 120 hours, and if a variant overlapped with a reported peak in any of the three replicates (GSM2083802, GSM2083803, GSM2083804) (52), it was denoted as overlapping.

### Percentage of True Causal Variants in HL-60 Undifferentiated Cells

In HL-60 undifferentiated cells, our called positive rate was 23.8% given that 99 of 416 candidates were significant (p-value < 0.05). The sensitivity of this assay is determined through the number of times eCROPseq previously validated positive controls^25^ (rs2251039 for *CISD1* and rs35675666 for *PARK7*) were significant in each of the 12 transductions. In total, we observed a 62.5% sensitivity rate as 15 out of 24 times the positive controls were significant. Specificity of the assay can be determined from one minus the false positive rate determined by the percentage of negative controls that were significant. In undifferentiated HL-60 cells, the false positive rate was 7.4%, thus the specificity was 92.6%. Using this called positive rate, sensitivity, and specificity, results in an estimation that only 30% of the variants are true positives, at a precision of 78% and overall accuracy of 0.84.

However, knowing positive controls are likely to have larger effects and that our assay included some mislabeled variants as candidates from false positives in our initial screening, we can reduce the sensitivity of the positive controls to be more reflective of the candidates tested and percentage of true positives to be closer to 50%. Specifically, a sensitivity of 40.5% maintains a high positive predictive value (85%), negative predictive value (61%), and overall accuracy (0.67) and predicts 84 of the estimated 208 true causal variants are correctly identified while 124 are missed.

### Overlapping Significant and True Causal Candidates between Cell Types

Variants significant (p-value < 0.05 with a minimum of five cells) across all three cell types were compared, resulting in 18 overlaps between HL-60 undifferentiated and macrophage, 14 between undifferentiated and neutrophil, and 16 between macrophage and neutrophil. We then predicted the number of true causal candidates across any pairwise comparison. Following our approach in Percentage of True Causal Variants in HL-60 Undifferentiated Cells, we assume a 50% true positive rate and utilize the known specificity, sample size of candidates detected in both cell types, and overall significance rate of candidates (tables S2-S4) to estimate sensitivity levels between 33-45% for any cell type. For example, out of the 322 variants tested in both undifferentiated HL-60 and macrophages, we determined 45% sensitivity for undifferentiated HL-60 and 33% for macrophage, which yields 161 true causal candidates of which 72 are correctly identified in undifferentiated HL 60 and 53 in macrophage (table S2). Across any pairwise comparison, the positive predictive value remains greater than or equal to 0.82 and negative predictive value (NPV) greater than or equal to 0.58 with overall accuracy never below 0.63.

We then calculate the number of expected overlapping true causal variants in a more highly powered assay by dividing the number of overlapping significant candidates (18 in undifferentiated HL-60 and macrophage, 14 in undifferentiated HL-60 and neutrophil, and 16 in macrophage and neutrophil) by the multiplied sensitivities of each cell type to infer a proportion of overlap between each cell type (75% between undifferentiated and macrophage, 54% between undifferentiated and neutrophil, and 87% macrophage and neutrophil) (Table 2).

### Simulations

To determine if simulated effects of our detected causal candidates can explain as much variance as an effect by the lead variant of the peak, we first selected a target gene with two detected eQTL candidates. Specifically, for *NFATC1*, we detected rs9748916 to be significant (p-value < 0.05) in macrophages and rs4799052 in neutrophils. The lead eQTL variant, rs9962906, was tested in our assay but was not significant. We then downloaded 1000 Genomes Phase 3 whole genotype data for 2,504 individuals^62^ for each variant from (https://jan2020.archive.ensembl.org/). We denoted the number of ancestral alleles reported for an individual for each variant (zero, one, or two). We then created a baseline expression level for each individual by utilizing a random normal distribution in Rstudio using rnorm(). To simulate gene expression of our lead variant model, we multiplied each baseline by the product of the number of lead variant ancestral alleles (LV) and effect size (β). Thus, simulated gene expression = baseline x (LV x β). To simulate gene expression of our significant candidate model, we multiplied each baseline by the product of the number of candidate 1 ancestral alleles (Var1) and effect size (β1) as well as the product of the number of candidate 2 ancestral alleles (Var2) and effect size (β 2). Thus, simulated gene expression = baseline x (Var1 x β1) x (Var2 x β2). We determined the amount of variance explained by either the effects of two candidates or an effect by the lead variant by determining the R2 of each respective linear model in Rstudio: lm(simulated gene expression ∼ lead variant genotype).

Utilizing an effect size of 0.5 for rs9748916 and −0.4 for rs4799052 and an effect size in between the two of 0.45 for the lead variant, our two-candidate model explained 6.5% of variance whereas the lead variant model explained 6.1%, as determined by the *R*^2^ value. Modeling with different effect sizes consistently shows situations in which the candidates can explain just as much or even more variance than the lead variant.

The second lead variant did not have whole genotype data available, so we repeated this analysis with the third most significant variant in the peak, rs9748849. Similarly to the lead variant, rs9748849 was tested in our eCROPseq assay but was not significant. Using the same baseline gene expression and multiplying by the product of the number of ancestral alleles present by effect size of 0.45, only 5.4% of variance was explained, indicating that not only can our candidates explain the major eQTL signal, but also other variants in the peak. Similar results are readily obtained across a range of estimated effect sizes, confirming prior computational results implying that the lead variant often captures the effects of two or more adjacent variants in high linkage disequilibrium^40^.

### Adenine (A) to guanine (G) base editing for selected variants through lentivirus

To validate the significant variants identified from eCROPseq assay in base editing, variant candidates from the *DAP* and *CARS* genes were screened for A to G conversion. Lentiviral vector, pRDA_479^63^ (Addgene, 179099) was chosen, which contains an ABE8e (SpG) to enable NGN-PAMs targeting, a gRNA scaffold to produce gRNA and a puromycin selection marker. The gRNAs were designed to avoid NGG-PAMs, targeted by Cas9 nuclease from the HL-60/S4-Cas9 cell line. As a result, we selected two validated significant candidates of gene *DAP* (rs2964790 and rs2918392), another candidate that validated during analysis with all cells but not in analyses that removed cells without a detectable gRNA (rs10866479), as well as a previously not significant negative control (rs12654966). For *CARS*, three validated candidates (rs10488673, rs7929049, and rs440130) and one negative control (rs756693) were selected. Two previously used non-targeting gRNAs were also included as negative controls and results for all samples are visualized in Fig. 8. Guide RNA plasmids were prepared individually for each variant^64^. In detail, gRNA oligos were acquired from Eurofins Genomics and annealed using T4 Polynucleotide Kinase (NEB, M0201L).

Ligation of gRNA and digested vector (13,642 bp) was reacted by Quick Ligation kit (NEB, M2200L). The ligation product was chemically transformed into DH10B Competent Cells (Thermo Scientific, EC0113), incubating for 16-18 hours at 37°C. Single colonies were then picked up, cultured, mini prepped, and validated by Sanger sequencing.

Lentivirus was produced and measured similarly to the methods described for the other experiments. Spinfection was performed with a wider MOIs range (0.1-10). Transfected cells were then under 0.4 µg/ml puromycin selection for 8 days, followed by 10 days of recovery in a regular medium. Afterward, gDNAs from the cells were extracted (DNeasy Blood and Tissue Kits for DNA Isolation, Qiagen, 69504) to validate the editing efficiency by Sanger sequencing and ICE analysis. RNAs were extracted, and converted to cDNA by reverse transcription, for gene expression quantification by qRT-PCR. Two housekeeping genes, beta-actin and 18s rRNA were used to normalize qPCR outcomes.

## Acknowledgments

We are grateful to many individuals for discussion and encouragement and particularly note contributions from Andy Bass, Dave Cutler, Maggie Brown, John Rioux and Judy Cho, as well as administrative support from Stefanie Boettle.

## Author contributions

Conceptualization and funding acquisition: GG, GB. Investigation: MC, EG. Supervision: CL, GB, GG. Experimental support: BM, AL. Analysis: EG. Writing – original draft: EG, MC, GG. Writing – review and editing: EG, MC, CL, GB, GG.

## Data and materials availability

All data, code, and materials used in the eQTL analysis are available at https://github.com/GibsonLab-GT/All-but-One and can be viewed interactively at https://eqtlhub-gt.shinyapps.io/shiny/. All guide RNAs, lists of SNPs, and analytical results are itemized in the supplementary tables.

## Supplementary Materials

Available from Corresponding Author Greg Gibson upon request

Figs. S1 to S9

Tables S1 to S10

